# Early midbrain sensorimotor pathway is involved in the timely initiation and direction of swimming in the hatchling *Xenopus laevis* tadpole

**DOI:** 10.1101/2022.08.24.505113

**Authors:** Michelle Christine Larbi, Giulia Messa, Helin Jalal, Stella Koutsikou

## Abstract

Vertebrate locomotion is heavily dependent on descending control originating in the midbrain and subsequently influencing central pattern generators in the spinal cord. However, the midbrain neuronal circuitry and its connections with other brainstem and spinal motor circuits has not been fully elucidated. Basal vertebrates with very simple nervous system, like the hatchling *Xenopus laevis* tadpole, have been instrumental in unravelling fundamental principles of locomotion and its suspraspinal control. Here, we use behavioral and electrophysiological approaches in combination with lesions of the midbrain to investigate its contribution to the initiation and control of the tadpole swimming in response to trunk skin stimulation. None of the midbrain lesions studied here blocked the tadpole’s sustained swim behavior following trunk skin stimulation. However, we identified that distinct midbrain lesions led to significant changes in the latency and trajectory of swimming. These changes could partly be explained by the increase in synchronous muscle contractions on the opposite sides of the tadpole’s body and permanent deflection of the tail from its normal position, respectively. Furthermore, the midbrain lesions led to significant changes in the tadpole’s ability to stop swimming when it bumps head on to solid objects. We conclude that the tadpole’s embryonic trunk skin sensorimotor pathway involves the midbrain, which harbors essential neuronal circuitry to significantly contribute to the appropriate, timely and coordinated selection and execution of locomotion, imperative to the animal’s survival.

## Introduction

Animal survival relies heavily on the selection and execution of appropriate and well-timed motor actions. These could include walking, swimming, maintaining balance, moving of the eyes, running away from a predator, breathing, chewing, reaching out for objects and many more depending on the species and its environment (Orlovsky et al., 1999). In vertebrates, an important role of the central nervous system (CNS) is the generation and execution of locomotion, one of the most fundamental and extensively studied motor actions that enables full body propulsion (Grillner and El Manira, 2020).

The spinal central pattern generator (CPG) circuits are responsible for the generation of the basic locomotor rhythm, by establishing the appropriate sequence of muscle activation combined with reciprocal muscle inhibition (Buchanan and Grillner, 1987;Kiehn, 2006;Goulding, 2009;Roberts et al., 2010;Kiehn, 2016). In order to meet everchanging behavioral demands, the activity of spinal CPG is modulated by descending inputs originating in highly distributed supraspinal neuronal circuits (Dubuc et al., 2008;Lemon, 2008;El Manira and Grillner, 2014;Bouvier et al., 2015;Daghfous et al., 2016;Hsu et al., 2017;Arber and Costa, 2018;Ferreira-Pinto et al., 2018;Ruder and Arber, 2019;Grillner and El Manira, 2020;Arber and Costa, 2022).

The midbrain is an integral part of the survival brain network, and its role has been found to influence behavioral motor actions by projecting onto other parts of the brainstem, which in turn modulate CPG circuit activity in the spinal cord (Koutsikou et al., 2014;Koutsikou et al., 2015;Koutsikou et al., 2017;Arber and Costa, 2018;Ferreira-Pinto et al., 2018;Grillner and El Manira, 2020;Arber and Costa, 2022). In particular, the locomotor command systems in the midbrain (MLR: mesencephalic locomotor region), initially identified in cats (Shik et al., 1966;Shik et al., 1969), are conserved within the vertebrate lineage and play a multifaceted role in the control of locomotion (Ryczko and Dubuc, 2013;Grillner and El Manira, 2020;Carbo-Tano et al., 2022). Additionally, the descending circuitry of the midbrain nucleus of the medial longitudinal fascicle (nMLF) is the origin of the commands that regulate steering and posture during locomotion in zebrafish larvae (Severi et al., 2014;Thiele et al., 2014;Wang and McLean, 2014).

As in all vertebrates, supraspinal activity in the hatchling *Xenopus laevis* tadpole is present at initiation and during swimming (Roberts et al., 2010;Buhl et al., 2012;Koutsikou et al., 2018;Roberts et al., 2019). A dedicated descending pathway via the tadpole’s midbrain is responsible for sensory activation and modulation of swimming following light dimming (Roberts, 1978;Foster and Roberts, 1982;Jamieson and Roberts, 1999;2000). More specifically, a decrease in light intensity (light dimming) leads to excitation of the pineal eye photoreceptors, which in turn activate pineal ganglion cells with bilateral axons that could contact the diencephalic / mesencephalic descending (D/MD) neurons located in the midbrain. Axons of the D/MD neurons project ipsilaterally to the hindbrain where they could excite the CPG and mediate or modulate the swim response (Jamieson and Roberts, 1999).

Anatomical evidence indicates that the tadpole’s midbrain might also be involved in the initiation of swimming in response to trunk skin stimulation. Axons of trunk skin sensory Rohon-Beard neurons contact sensory pathway neurons, dla (dorsolateral ascending) and dlc (dorsolateral commissural), whose axons project to both sides of the hindbrain (Roberts and Clarke, 1982;Sillar and Roberts, 1988;1992;Li et al., 2001). However, most dla and some dlc axons also project to midbrain (Li et al., 2001). We understand that dla and dlc sensory pathway neurons initiate or accelerate swimming by amplifying excitation (Li et al., 2001;Li et al., 2004b). However, the function of the dla and dlc ascending projections to the midbrain is currently unclear. The presence of these projections suggests that *(i)* the midbrain might be essential part of the trunk skin sensorimotor pathway, even at this early stage of development and *(ii)* there could be transitory as well as longer-lasting influences of midbrain neurons on hindbrain- and/or spinal cord-driven motor activity.

Although research has shown that the midbrain is an integral part of the supraspinal motor control network, from basal vertebrates to mammals, its underlying neuronal circuitry, and interactions with the rest of the brainstem and spinal cord are not fully elucidated. Here we present the first investigation on the role of midbrain in the initiation and maintenance of trunk skin-evoked swimming in the young *Xenopus laevis* tadpole. We use touch-evoked or electrical trunk skin stimulation and midbrain lesions to assess the importance of midbrain in mediating and/or modulating the swimming of the hatchling tadpole. We present evidence for changes in latency to initiation, trajectory and stopping of swimming as well as loss of tail postural control following midbrain lesions.

## Materials and Methods

### Animal ethics and surgery

All experimental procedures were carried out under the relevant guidelines and approved by the University of Kent Animal Welfare and Ethical Review Body (AWERB). *Xenopus laevis* embryos were supplied by the European *Xenopus* Resource Centre (EXRC; Portsmouth, UK) and kept at 16°C in tap water treated with commercially available aquarium water conditioner. All experiments were carried out on embryos at developmental stage 37/38 (Nieuwkoop and Faber, 1956), at room temperature (RT: 19°C).

All surgical procedures on the tadpoles were performed in a small custom-made dish filled with saline (NaCl 115mM, KCl 3mM, CaCl_2_ 2mM, NaHCO_3_ 2.4mM, MgCl_2_ 1mM, HEPES 10mM; pH 7.4). Tadpoles were briefly anesthetised in 0.1% MS-222 (Ethyl 3-aminobenzoate methanesulfonate; Sigma-Aldrich, UK) and pinned to a small rotating Sylgard block. Dissections were performed by hand with sharpened custom-made tungsten needles under a dissection microscope. Figure 1A, B shows the area of the tadpole’s CNS exposed in preparation for the procedures described in detail below.

**FIGURE 1.**
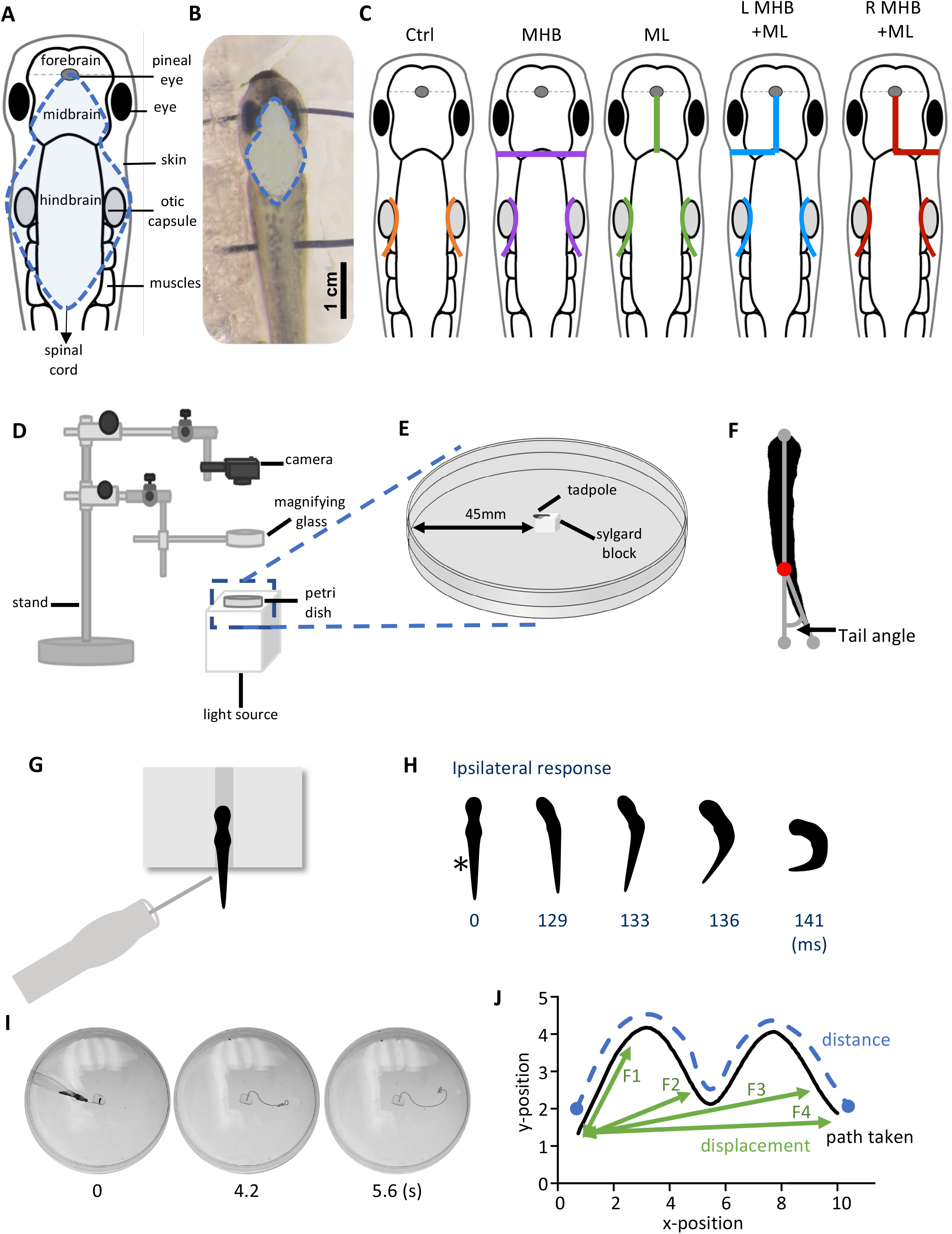
Midbrain lesions and design of behavioral experiments. Dorsal view of the *Xenopus laevis* tadpole at developmental stage 37/38. **(A)** Diagram of the tadpole’s anatomical features and key CNS regions. Blue dotted outline represents the area where the skin was opened to allow access to the brain. **(B)** The tadpole viewed under a dissection microscope with pins running across the body, through the eyes and the notochord just caudal to the obex. Blue dotted line denotes the area of open skin with the brain visible. **(C)** Representation of the midbrain lesions. Ctrl (Control, orange); trigeminal nerve transection at the level of the otic capsule. The trigeminal nerve transections were performed on all experimenal groups. MHB (Midbrain-Hindbrain Border lesion, purple); a transverse lesion through the MHB. ML (Midline lesion, green); lesion along the midline of the midbrain. L MHB + ML (Left MHB and ML lesion, blue); combination of left MHB and ML lesions. R MHB + ML (Right MHB and ML lesion, red); combination of right MHB and ML lesions. **(D)** Schematic diagram of the behavioural set up with **(E)** a zoomed in section of the tadpole’s starting position (not drawn to scale). **(F)** Illustration of the tail angle measurement used in subsequent figures. Red dot indicates the level of the anus, referred to as the starting point of the tail **(G)** The tadpoles starting position from a dorsal view. The tadpole is positioned upright into a groove carved onto a Sylgard block. Stimulation is applied on the left side of the tadpole’s trunk skin, at the level of the anus, using a glass pipette with an attached fine rabbit hair (custom-made). **(H)** Individual frames taken from high-speed videos recorded at 420 frames per second (fps). A short poke with a single rabbit hair to trunk skin receptors on the left side (*; time = 0 ms) initiated a swim response at 129 ms. **(I)** Individual video frames of a tadpole’s swim trajectory captured from a 30fps recording. In this example the control animal trial starts at stimulation (0 s) to the end of the tadpole’s swim cycle (5.6 s). A novel tracking software, FastTrack Software, was used to detect and trace the tadpole’s body using an automatic tracking algorithm. **(J)** Schematic showing the path taken (black), the distance (blue) and the displacement (green). Here, the displacement is described as the shortest distance from the starting position to the next frame (position), with frame intervals indicated as F1, F2 and F3, with F4 representing the final displacement. The distance here is described as the measure of the path taken from the starting to the final position. Abbreviations: Ctrl, control; MHB, midbrain-hindbrain border; ML, midbrain midline; L MHB + ML, left midbrain-hindbrain border and midbrain midline; R MHB + ML, right midbrain-hindbrain border, and midbrain midline.

Both trigeminal nerves were severed at the level of the otic capsule (Figures 1A, C), to block skin impulses entering the nervous system and initiating swimming (Roberts, 1996;James, 2009;James and Soffe, 2011). The contribution of the hatchling tadpole’s midbrain to the control of movement was assessed through lesions as depicted in Figure 1C. These included: *(i)* a transverse lesion along the midbrain-hindbrain border (MHB), *(ii)* lesion along the midbrain’s midline (ML), *(iii)* lesion along the midbrain’s midline in combination with a transverse lesion on the left (L MHB +ML), and *(iv)* the right (R MHB +ML) side of the MHB.

### Behavior

Following the above preparatory procedures, all animals were allowed to recover in saline for 10 minutes at RT. Behavioral experiments were carried out in a Petri dish (diameter 90 mm) filled with saline, which was shallow enough to prevent animals from swimming in an upward spiral (Jamieson and Roberts, 2000;Roberts et al., 2000). The behavioral setup used is illustrated in Figure 1D, E. Briefly, animals were positioned dorsal side up within a groove made into a Sylgard block, which was affixed in the centre of the Petri dish. The Petri dish was illuminated from below by an LED light to enhance the contrast of the tadpole’s silhouette during video recording. A digital camera (Exilim EX-FH100, Casio) was positioned above this setup and was used to record high-speed videos in black and white mode. Videos were recorded at 420 fps for experiments in which latency to swim initiation was studied. Additionally, 30 fps videos were recorded in experiments where the swimming trajectory was analysed.

The angle of the tail was measured before swimming (at rest). Using the ImageJ software (NIH), we determined the tail angle based on the deviation of the tail from a straight line running from the front of the tadpole’s head (gray dot; Figure 1F) through the beginning of the tail (red dot; Figure 1F) to the most caudal tip of the tail (gray dot; Figure 1F).

A short poke with a fine (rabbit) hair was manually delivered to the tadpole’s trunk skin on the left side of its body to initiate swimming (Figure 1G). Each animal was allowed recovery time of 2 minutes between trials. Latency videos were analysed using the ImageJ software. The latency (ms) to swim initiation was calculated based on the number of video frames from trunk skin stimulation (time 0 ms on Figure 1H) to the first head bend (time 129 ms on Figure 1H). The direction of head movement was also recorded as either on the ipsilateral or contralateral side in relation to the stimulus (ipsilateral to the stimulus on Figure 1H).

FastTrack Software, an open-source tracking software (Gallois and Candelier, 2021), was used to detect and track the tadpole in experiments on swimming trajectory (Figure 1I). A text file containing the X-Y position of the tadpole’s body across frames was generated and kinematic analysis was preformed using MS Excel.

The following equation was used to calculate the displacement:

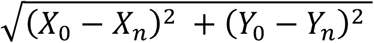

whereby, X_0_ and Y_0_ is the starting position of the tadpole and n is the position of the tadpole in the frame under analysis (schematic illustration shown in Figure 1J).

### Electrophysiology

Preparatory procedures for the electrophysiological recordings of fictive swimming also included removal of the skin covering the trunk muscles on both sides of the tadpole’s body for access to the myotomal clefts. After surgery, animals were allowed to recover and tested for robust swimming prior to being paralyzed in α-bungarotoxin (0.01M, Invitrogen) for 50 minutes at RT. Animals were pinned on a rotating Sylgard block in a small recording dish filled with saline (Figure 1B). Two borosilicate glass suction electrodes (diameter ∼50μm) were attached to both sides of the tadpole’s body approximately at the level of the 4^th^ cleft to record ventral root (VR) activity (Figure 2A, B). Such positioning of the VR recording electrodes permits accurate capturing of the side of fictive swimming initiation. A third glass suction electrode was attached to the trunk skin on the right side of the body, at the level of the anus, to deliver electrical stimuli to the trunk skin. A schematic view of the electrode positions can be seen in Figures 2A, B (Koutsikou et al., 2018;Messa and Koutsikou, 2021). Electrical stimulation was delivered via a custom-made TTL pulse generator automatically driven through software (Signal 7, CED, Cambridge, UK). Threshold stimulation was set as the smallest stimulus (both in intensity -V- and duration -ms-) that led to swim initiation. Suprathreshold stimulation was defined as the intensity of threshold stimulus + 1V, with the same duration as of threshold stimulus. Threshold stimulation was verified in each animal prior to experimental recordings. Subsequently, suprathreshold stimulus was calculated for every animal accordingly. All animals started swimming with a threshold stimulus in the range of 3.5-4.5 V and 0.25-0.4 ms. The electrical signal from left and right VR was amplified, 50/60Hz noise was eliminated via a noise eliminator (HumBug, Digitimer, UK), and data were acquired through Power 1401 mkII (CED, Cambridge, UK) in Signal 7 (CED, Cambridge, UK) and displayed as shown in Figures 2C, D.

**FIGURE 2.**
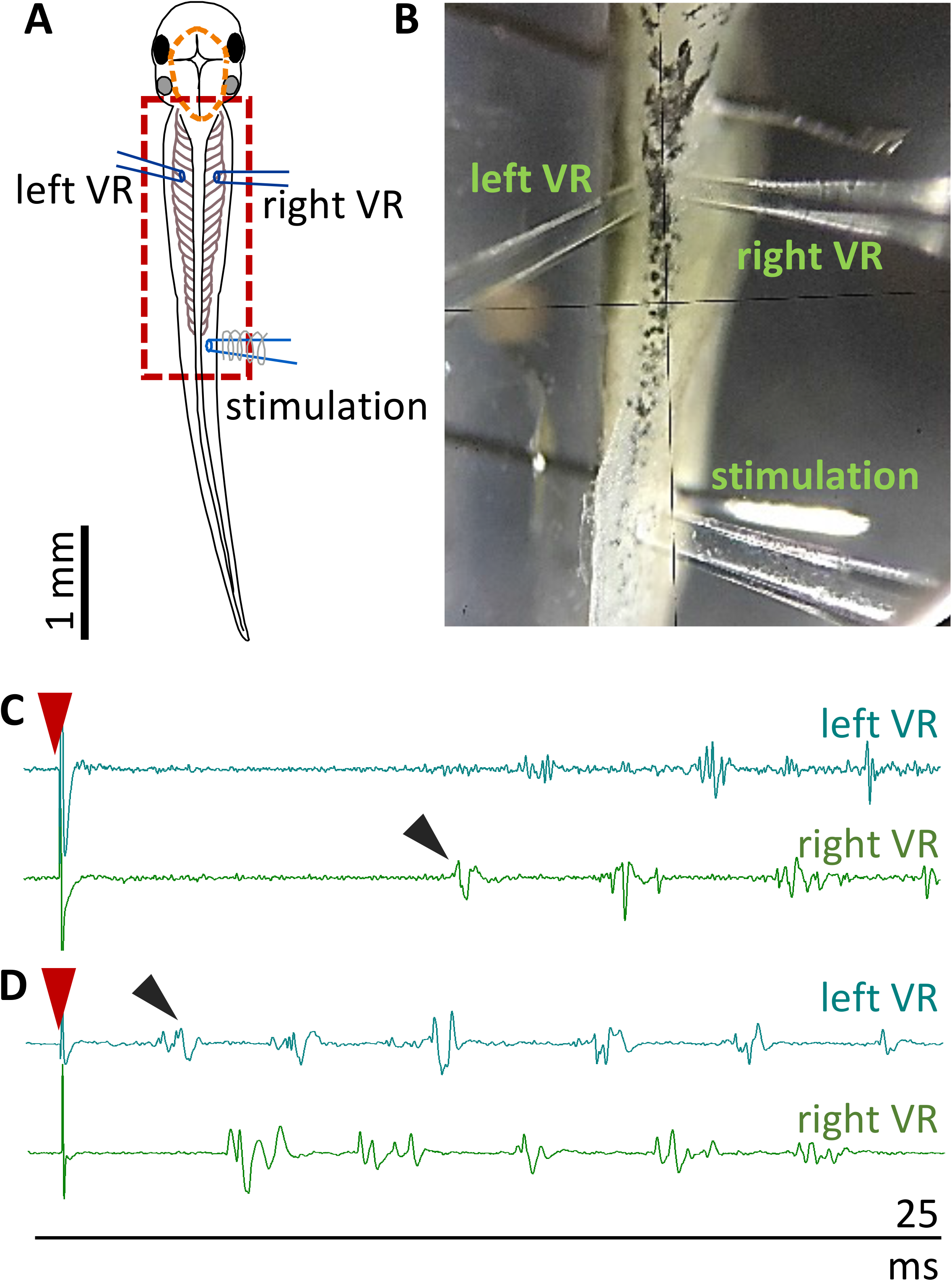
Experimental setup for fictive swimming. **(A)** Dorsal view of the tadpole with recording and stimulating electrodes in postition. Left and right VR (ventral root) electrodes were positioned facing each other approximately at the 4th myotomal cleft. The stimulating electrode was always positioned on the right side at the level of the anus. Orange dashed line denotes the area of open skin with the brain visible. Red dashed square area indicates the area presented in panel B. **(B)** Picture of the trunk of a tadpole pinned to the Sylgard block during extracellular VR recordings. Right and left VR and stimulating electrodes, are positioned as described in A. **(C)** Example of an ipsilateral start of fictive swimming in response to electrical trunk skin stimulation (red arrowhead). The first VR burst is recorded on the right side of the body (right VR, ipsilateral; black arrowhead). **(D)** Example of a contralateral start of fictive swimming in response to electrical trunk skin stimulation (red arrowhead). The first VR burst is recorded on the left side of the body (contralateral, left VR; black arrowhead). In all experiments stimulation was delivered on the right side of the tadpole’s body.

### Statistical Analysis

All statistical analyses were performed using Prism 9 (GraphPad). Data were tested for normality using Shapiro-Wilk test, with normality criterion set at p<0.05. The statistical tests are as stated throughout the Results. To describe the central tendency and variability, median with interquartile range values (IQR: 25-75 percentile) were used unless stated otherwise. Statistical analysis was not performed on percentages.

## Results

None of the lesions described here abolished the swim response. This agrees with previous findings showing that the tadpole can generate episodes of sustained swimming after removal of the CNS rostral to the seventh rhombomere (Li et al., 2006). However, the specific consequences of such lesions have not previously been examined in the hatchling tadpole.

### The midbrain contributes to the latency and side of first motor response in freely moving animals

Following a gentle touch to the tadpole’s trunk skin, swimming is initiated. With the use of high-speed videos recorded at 420 fps, the latency (ms) from the touch stimulus to swim initiation was measured (Figure 3A). The latency to swim initiation measured in MHB-lesioned animals (n=14 tadpoles) was significantly longer compared to controls (n=10 control tadpoles; p<0.0001, Kruskal-Wallis/Dunn’s test; control group median: 104.8, IQR: 73.2-128.0 ms *vs* MHB-lesioned median: 156.0, IQR: 125.0-243.5 ms). Tadpoles with the R MHB +ML lesion (n=9 tadpoles) also showed significantly longer latencies compared to control animals (n=10 control tadpoles; p=0.0011, Kruskal-Wallis/Dunn’s test; control vs R MHB +ML median: 126.2, IQR: 104.8-198.8 ms; Figure 3A).

**FIGURE 3.**
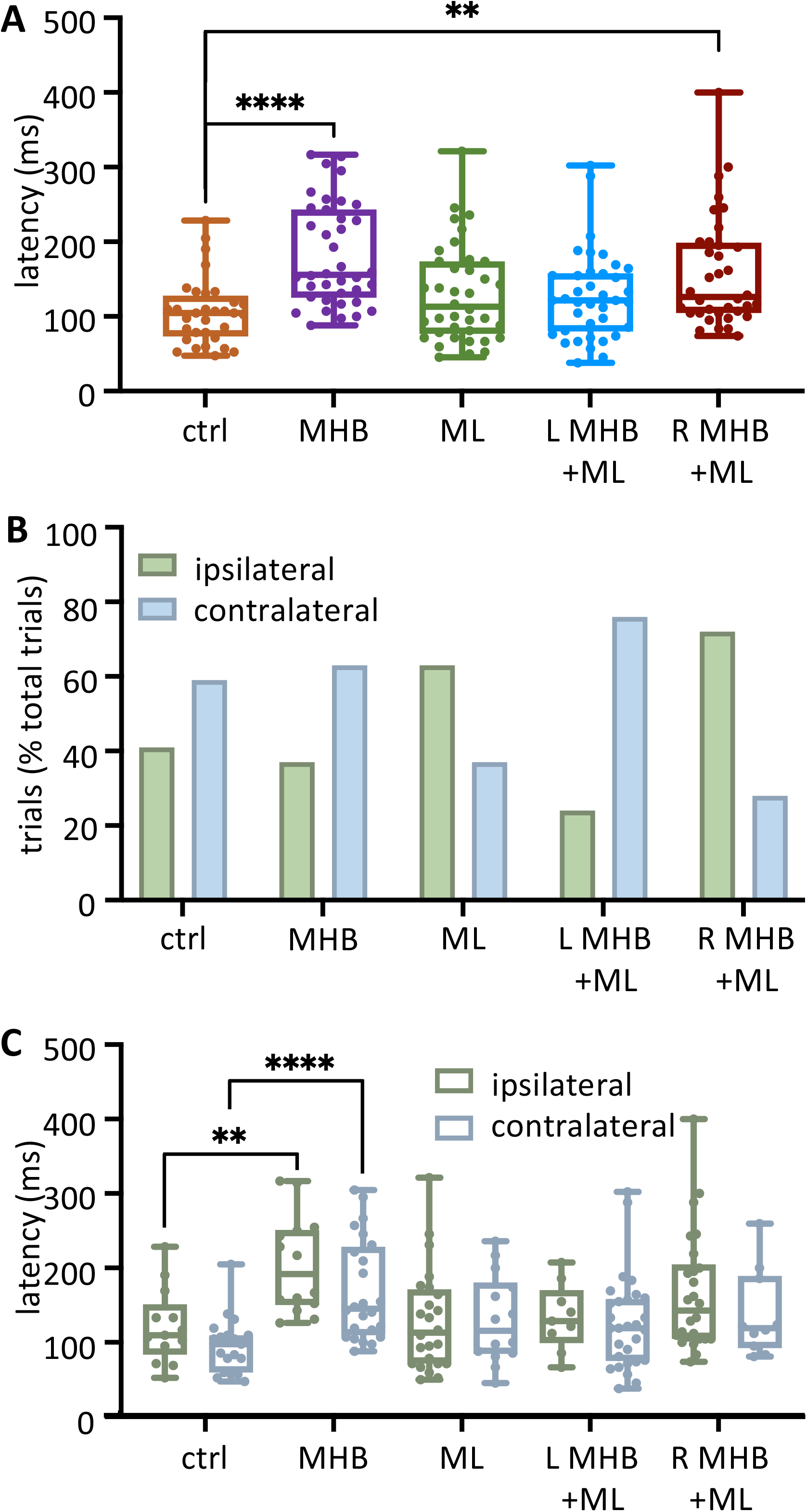
Midbrain lesions lead to changes in latency and side of swim initiation in freely moving animals. **(A)** Latency (ms) to the start of swimming in response to trunk skin stimulation. Ctrl: 104.8, 73.2-128.0 ms, MHB: 156.0, 125.9-243.5 ms, midline: 113.1, 76.79-173.8 ms, L MHB+ML: 121.4, 79.76-157.7 ms, R MHB+ML: 126.2, 104.8-198.8 ms. **(B)** Percentage occurrence (% total number of trials for each experimental group) of the first behavioral motor response in relation to the side of the stimulus. Ctrl: 41% ipsilateral, 59% contralateral; MHB: 37% ipsilateral, 63% contralateral; Midline: 63% ipsilateral, 37% contralateral; L MHB+ML: 24% ipsilateral, 76% contralateral; R MHB+ML: 72% ipsilateral, 28% contralateral. **(C)** Latency to the initiation of swimming in ipsilateral *vs* contralateral starts. Ctrl ipsilateral 109.5, 83.3-151.2 ms, contralateral 97.6, 59.5-109.5 ms; MHB ipsilateral 191.7,150.0-251.2 ms, contralateral 145.2, 109.5-228.6 ms; Midline ipsilateral 113.1, 71.4-171.4 ms, contralateral 115.5, 84.5-180.4 ms; L MHB+ML ipsilateral 128.6, 98.8-170.2 ms, contralateral 119.1, 75.0-158.3 ms; R MHB+ML ipsilateral 142.9, 104.8-204.8 ms, contralateral 119.1, 92.3-189.3 ms. For all panels, all data reported as median, 25-75 percentile. Ctrl: n=10 tadpoles, trials=32; MHB: n=14 tadpoles, trials=38; Midline: n=11 tadpoles, trials=38; L MHB+ML: n=12 tadpoles, trials=38; R MHB+ML: n=9 tadpoles, trials=36. For boxplot (A,C) single data points are plotted, the middle horizontal line represents median latency, the box represents the interquartile range (IQR, 25-75 percentile), and the error bars represent the minimum and maximum values. Results are reported as median and IQR. ****p<0.0001, **p<0.01 for Kruskal-Wallis/Dunn’s test.

Excitability due to sensory stimulation on one side of the skin is transmitted to both sides of the hindbrain and can initiate swimming on either side of the tadpole’s body (Buhl et al., 2012;Buhl et al., 2015;Roberts et al., 2019;Ferrario et al., 2021;Messa and Koutsikou, 2021). In this study, following left trunk skin stimulation, control tadpoles (n=10) initiated swimming more often on the unstimulated side (contralateral to the stimulus; Figure 3B) at percentages of 41% ipsilateral (13/32 trials) and 59% contralateral (19/32 trials). This relationship between ipsilateral and contralateral swim initiations has been affected, to varying degrees, by all midbrain lesions. More specifically, MHB-lesioned animals (n=14) showed a slight increase in the percentage of contralateral starts (37% ipsilateral 14/38 trials *vs* 63% contralateral 24/38 trials). Interestingly, the partial lesion to the MHB in combination with the ML lesion (L MHB + ML; n=12 tadpoles; Figure 2B) increased even further the percentage of contralateral swim starts (24% ipsilateral 9/38 trials *vs* 76% contralateral 29/38 trials). In contrast to these lesions favoring contralateral starts, the ML (n=11 tadpoles; 63% ipsilateral, 24/38 trials *vs* 37% contralateral, 14/38 trials) and the R MHB + ML lesions (n=9 tadpoles; 72% ipsilateral 26/36 trials *vs* 28% contralateral 10/36 trials) led animals to more frequent initiation of swimming on the ipsilateral side (Figure 3B).

The latency data were further analysed in relation to the side of swim start (Figure 3C; number of tadpoles as mentioned above). Within each tadpole group, there was no significant difference in the initiation latency between ipsilateral *vs* contralateral starts (p>0.05, individual Mann-Whitney test within each tadpole group comparing ipsilateral *vs* contralateral latency: ms; Figure 3C). However, comparisons across groups revealed that the ipsilateral and contralateral latencies for only the MHB-lesioned animals differed significantly when compared to the respective values obtained from the control animals (Figure 3C; p=0.0011; control ipsilateral median: 109.5, IQR: 83.3-151.2 ms *vs* MHB ipsilateral median: 191.7, IQR: 150.0-251.2 ms; p<0.0001; control contralateral median: 97.6, IQR: 59.5-109.5 ms *vs* MHB contralateral median: 145.2, IQR: 109.5-228.6 ms; Mann-Whitney test). Overall, the complete disconnection of the midbrain (MHB) from the rest of the brainstem had the greatest effect on latency, indicating the involvement of the midbrain in the modulation of motor initiation. Furthermore, the change to the side of the initial body bend, for ML- and R MHB + ML-lesioned animals, suggests the importance of possible midbrain commissural connections (currently unknown) in determining the side of the first motor response.

### The midbrain contributes to the latency, side of first motor response and alternating initiation of fictive swimming

In the behavioral experiments above, the MHB lesion caused the most significant change in latency to initiation of swimming when compared to control animal responses. For this reason, we used the MHB-lesioned *vs* control tadpoles to further investigate the reasons behind this change to swim latency. In this set of experiments, with electrical stimulation delivered on trunk skin at threshold intensities, MHB-lesioned tadpoles (n=7) showed shorter latency to first ventral root (VR) activity in comparison to control animals (n=5 tadpoles; Figure 4A; Mann-Whitney test p=0.0049, control median: 100.9, IQR: 91.0-117.8 ms *vs* MHB lesioned median: 48.1, IQR: 36.2-61.4 ms;). We also identified the side of the first VR burst (Figure 2C, D). The control group activated the ipsilateral side first in 63.6% of the trials (14/22 trials), whilst the first VR burst appeared on the contralateral side in 36.4% of the trials (8/22 trials; Figure 4B). Additionally, it was revealed that tadpoles with lesioned MHB failed to start efficient swimming (with alternating VR activity) at the first VR burst in 13.8% of the trials (4/29 trials; Figure 4B). Indeed, in 4/29 trials VR bursts were detected simultaneously on both sides of the body of MHB-lesioned tadpoles (referred as ‘synchrony’ in Figure 4B and our previous work: (Ferrario et al., 2021)). The latency to fictive swimming initiation was then grouped and plotted based on the side of the first VR recorded (Figure 4C). Although no significant difference was identified (Figure 4C; control: p=0.365, Mann-Whitney and MHB-lesioned animals: p=0.1859, Kruskal-Wallis), the latencies for synchronous start in MHB-lesioned tadpoles appeared less variable than latencies reported for contralateral or ipsilateral initiation in both lesioned and control animals (Figure 4C; control ipsilateral median: 101.9, IQR: 93.1-112.0 ms; control contralateral median: 67.44, IQR: 23.6-132.8 ms; MHB ipsilateral median: 49.8, IQR: 39.6-64.0 ms; MHB contralateral median: 50.7, IQR: 36.5-83.6 ms; MHB synchrony median: 36.3, IQR: 35.9-39.8 ms).

**FIGURE 4.**
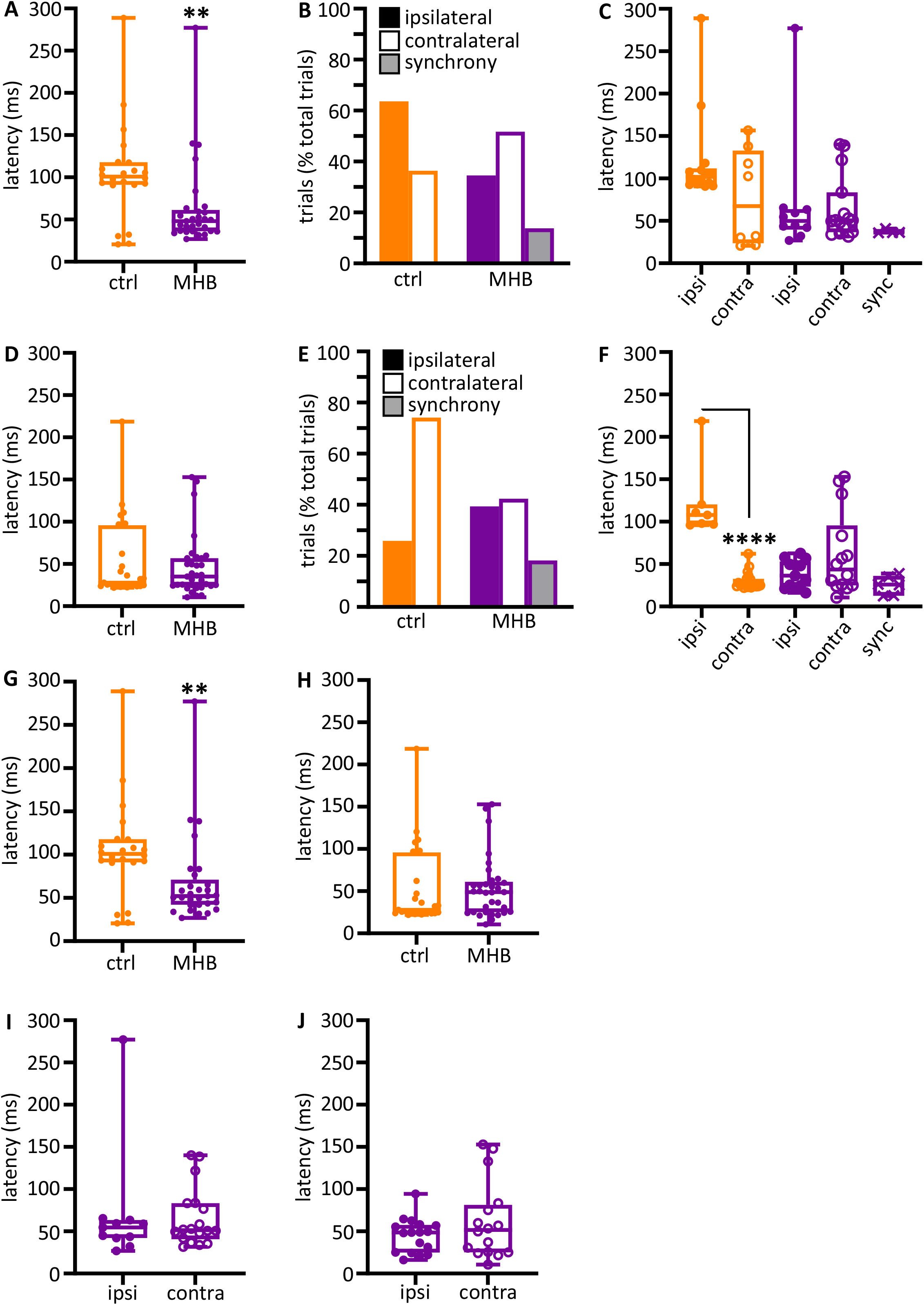
Midbrain-hindbrain border (MHB) lesion leads to changes in latency and side of fictive swim initiation. **(A)** Latencies (ms) to the first VR burst after threshold electrical stimulus was delivered to the trunk skin in control and MHB-lesioned animals. Mann-Whitney test *p*=0.0049, controls 100.9, 91.02-117.8 ms *vs* MHB lesioned 48.1, 36.24-61.4 ms; data reported as median, 25-75 percentile. Ctrl: n = 5 tadpoles, trials=22; MHB: n=7 tadpoles, trials=29. **(B)** Percentage occurrence (% total number of trials for each experimental group) of the first VR burst after threshold electrical stimulation in control (orange) and MHB-lesioned animals (violet). Ctrl ipsilateral 63.6% (14/22 trials), contralateral 36.4% (8/22 trials); MHB ipsilateral 34.5% (10/29 trials), contralateral 51.7% (15/29 trials), synchrony 13.8% (4/29 trials). **(C)** Latencies (ms) to the first VR burst after threshold electrical stimulus delivered to the trunk skin in control (orange) and MHB-lesioned (violet) animals. Solid circles represent latencies to ipsilateral first VR burst, open circles represent latencies to contralateral first VR burst, and crosses represent VR bursts recorded simultaneously on both sides of the body. Mann-Whitney test for the ctrl group, p=0.365; Kruskal-Wallis test for the MHB-lesioned group, p=0.1859. Ctrl ipsilateral 101.9, 93.1-112.0 ms, contralateral 67.4, 23.6-132.8 ms; MHB ipsilateral 49.8, 39.6-64.0 ms, contralateral 50.7, 36.5-83.6 ms, synchrony 36.2, 35.9-39.8 ms. Data are reported as median, 25-75 percentiles. Ctrl: n = 5 tadpoles, trials=22; MHB: n = 7, trials=29. **(D)** Latencies (ms) to the first VR burst after suprathreshold electrical stimulus delivered to the trunk skin in control and MHB-lesioned tapoles. Mann-Whitney test, p=0.8249; ctrl: 27.82, 24.35-95.81 ms; MHB: 35.16, 24.28-56.94 ms. Data reported as median, 25-75 percentile. Ctrl: n = 5 tadpoles, trials=27; MHB: n = 7, trials=33. **(E)** Percentage occurrence (% total number of trials for each experimental group) of the first VR burst after suprathreshold electrical stimulation of control (orange) and MHB-lesioned animals (violet). Ctrl ipsilateral 25.9% (20/27 trials), contralateral 74.1% (7/27 trials); MHB ipsilateral 39.4% (13/33 trials), contralateral 42.4% (14/33 trials), synchrony 18.2% (6/33 trials). **(F)** Latencies (ms) to the first VR burst after suprathreshold electrical stimulus delivered to the trunk skin of control (orange) and MHB-lesioned animals (violet). Solid circles represent latencies to ipsilateral first VR burst, open circles represent latencies to contralateral first VR burst, and crosses represent VR bursts recorded simultaneously on both sides of the body. Mann-Whitney test for the ctrl group, p<0.0001: ipsilateral 107.9, 97.0-120.3 ms, contralateral 25.5, 23.9-32.7 ms; Kruskal-Wallis test for MHB group, p=0.2108: ipsilateral 36.5, 23.3-53.2 ms, contralateral 43.6, 24.8-95.7 ms, synchrony 25.8, 12.7-36.1 ms. All data are reported as median, 25-75 percentile. Ctrl: n= 5 tadpoles, trials=27; MHB-lesioned: n = 7, trials=33). **(G)** Latencies (ms) to the first alternating VR burst (*i*.*e*., the first burst indicative of the start of fictive swimming) after a threshold electrical stimulus delivered to the trunk skin of control and MHB-lesioned tadpoles. Mann-Whitney test, p=0.0049; ctrl: 100.9, 91.0-117.8 ms; MHB: 52.1, 42.0-71.1 ms. Ctrl: n = 5 tadpoles, trials=22; MHB: n = 7, trials=29. **(H)** Latencies (ms) to the first alternating VR burst after a suprathreshold electrical stimulus delivered to the trunk skin of control and MHB-lesioned tadpoles. Mann-Whitney test, p=0.4879; ctrl: 27.8, 24.4-95.8; MHB: 48.9, 25.1-61.1 ms. Ctrl: n = 5 tadpoles, trials=27; MHB: n = 7, trials=33. **(I)** Latency (ms) for asynchronous starts (synchrony data are omitted) after a threshold stimulation and according to the side of the first VR burst (Mann-Whitney test, p=0.8079; ipsilateral 54.6, 42.1-63.5 ms; contralateral: 51.4, 40.6-83.5 ms; n = 7 tadpoles, trials=29). **(J)** Latency to asynchronous starts after a suprathreshold stimulation and according to the side of first VR burst (Mann-Whitney test, p=0.4654; ipsilateral 48.7, 24.8-57.6 ms, contralateral 51.6, 25.2-81.3 ms; n = 7 tadpoles, trials=33). In all panels, data collected on controls are in orange, data collected on MHB lesioned animals are in violet. In A, C, D, F, G-J single data points are plotted; boxes indicate 5-95 percentile; middle horizontal line in each box represents median value; error bars indicate minimum and maximum values. In panel B and E; filled bars: ipsilateral first VR burst; white bars: contralateral first VR burst; grey bars: synchronous VR first burst. All data reported in the figure legend are expressed as median, 25-75 percentile.

When electrical stimulation was delivered at suprathreshold intensities, latency to the first VR activity did not differ significantly in MHB-lesioned animals (n=7) compared to control group (n=5 tadpoles; Figure 4D; p=0.8249, Mann-Whitney test; control median: 27.8, IQR: 24.4-95.8; MHB median: 35.2, IQR: 24.3-56.9). In contrast to the response following threshold stimulation, control animals switched the side of first VR burst to the contralateral side (Figure 4E; contralateral 74.07% *vs* ipsilateral 25.93% (20/27 *vs* 7/27 trials respectively)). In MHB-lesioned tadpoles, the percentages of trials with ipsilateral *vs* contralateral initiation were similar following suprathreshold stimulation (ipsilateral 39.4% (13/33 trials) *vs* contralateral 42.4% (14/33 trials)). The percentage of synchronous initiations recorded was slightly higher in comparison to that recorded following threshold skin stimulation of MHB-lesioned animals (Figure 4E; 18.2% (6/33 trials) synchronous VR activity). At suprathreshold trunk skin stimulation, control tadpoles showed shorter latency to fictive swimming when started their movement on the contralateral side (Figure 4F; p<0.0001, Mann-Whitney test; ipsilateral median: 107.9, IQR: 97.0-120.3 ms *vs* contralateral median: 25.5, IQR: 23.9-32.7 ms).

Following MHB lesions, tadpoles failed to diversify their swim response. The latency to initiation of swimming was not significant across ipsilateral, contralateral, and synchronous fictive swim initiations (Figure 4F; p=0.2108, Kruskal-Wallis test; ipsilateral median: 36.5, IQR: 23.3-53.3 ms *vs* contralateral median: 43.6, IQR: 24.8-95.7 ms *vs* synchrony median: 25.8, IQR: 12.7-36.1 ms).

When VR activity appears synchronously on both sides of the tadpole’s trunk, the alternating contraction of antagonist muscles necessary for movement initiation cannot be achieved. The electrophysiological experiments here give us the opportunity to also identify and easily separate the latency to the first VR activity, post synchrony, which is the one leading to alternating fictive swimming. The latency of the first alternating VR was also measured and plotted for all MHB-lesioned animals *vs* the control group. At threshold stimulation, MHB-lesioned animals showed a significantly shorter latency compared to controls despite some of the alternating VR firing commencing post-synchrony (Figure 4G; p=0.0049, Mann-Whitney test; control median: 100.9, IQR: 91.0-117.8 ms; MHB lesion median: 52.2, IQR: 42.0-71.1 ms). On the contrary, at suprathreshold stimulation, latency values were not significant different between the two groups (Figure 4H; p=0.4879, Mann-Whitney test; control median: 27.8, IQR: 24.4-95.8 ms; MHB lesion median: 48.9, IQR: 25.1-61.1 ms).

When the latency values for alternating starts were plotted according to the side of initiation, MHB-lesioned tadpoles showed comparable latencies for both ipsilateral and contralateral fictive swim initiation following both threshold (Figure 4I; p=0.8079, Mann-Whitney test; ipsilateral median: 54.6, IQR: 42.1-63.5 ms; contralateral median: 51.4, IQR: 40.6-83.5 ms) and suprathreshold electrical trunk skin stimuli (Figure 4J; p=0.4654, Mann-Whitney test; ipsilateral median: 48.7, IQR: 24.8-57.6 ms; contralateral median: 51.6, IQR: 25.2-81.3 ms).

Both sets of experimental work described above, revealed that varying degrees of midbrain lesion led to changes in swim behavior. First, latencies to the initiation of swimming have been altered and it appears that the synchronous contraction of trunk muscles on opposite sides of the tadpole’s body might be in part responsible for such changes. Furthermore, we observed stark alterations to the side of swim initiation in midbrain-lesioned animals, in some instance leading to a reversal of the control behavior.

### Midbrain lesions affect the trajectory and maintenance of swimming in freely moving animals

The following experiments on freely moving animals focused on exploring the contribution and role of midbrain in the maintenance of trajectory and further aspects of sustained swimming (Figure 1I, J). Tadpole swim behavior was initiated in response to mechanical touch (fine rabbit hair) on the trunk skin (Figure 1G). The swim trajectories for each animal group were captured through video recording (Figure 1I) and subsequently traced, and plotted (Figures 1J, 5A, B). Representative examples from each experimental group are shown in Figure 5A (the trajectories of two animals per group are shown). The trajectories of control animals followed a forward direction, as the tadpoles swam away from the starting position (Figure 5A1). In contrast, the trajectories of midbrain-lesioned animals had a turning/circular pattern, pivoting around a specific area of the arena multiple times (often close to the starting point; Figure 5A2-5). Notably, R MHB + ML lesioned animals started briefly in a forward direction, which would turn into circular movement until the end of the swim episode (Figure 5A5). Further qualitative assessment showed that this group’s circular trajectories were also wider in diameter than the circular movements of all other midbrain-lesioned tadpoles (Figure 5A2-5).

**FIGURE 5.**
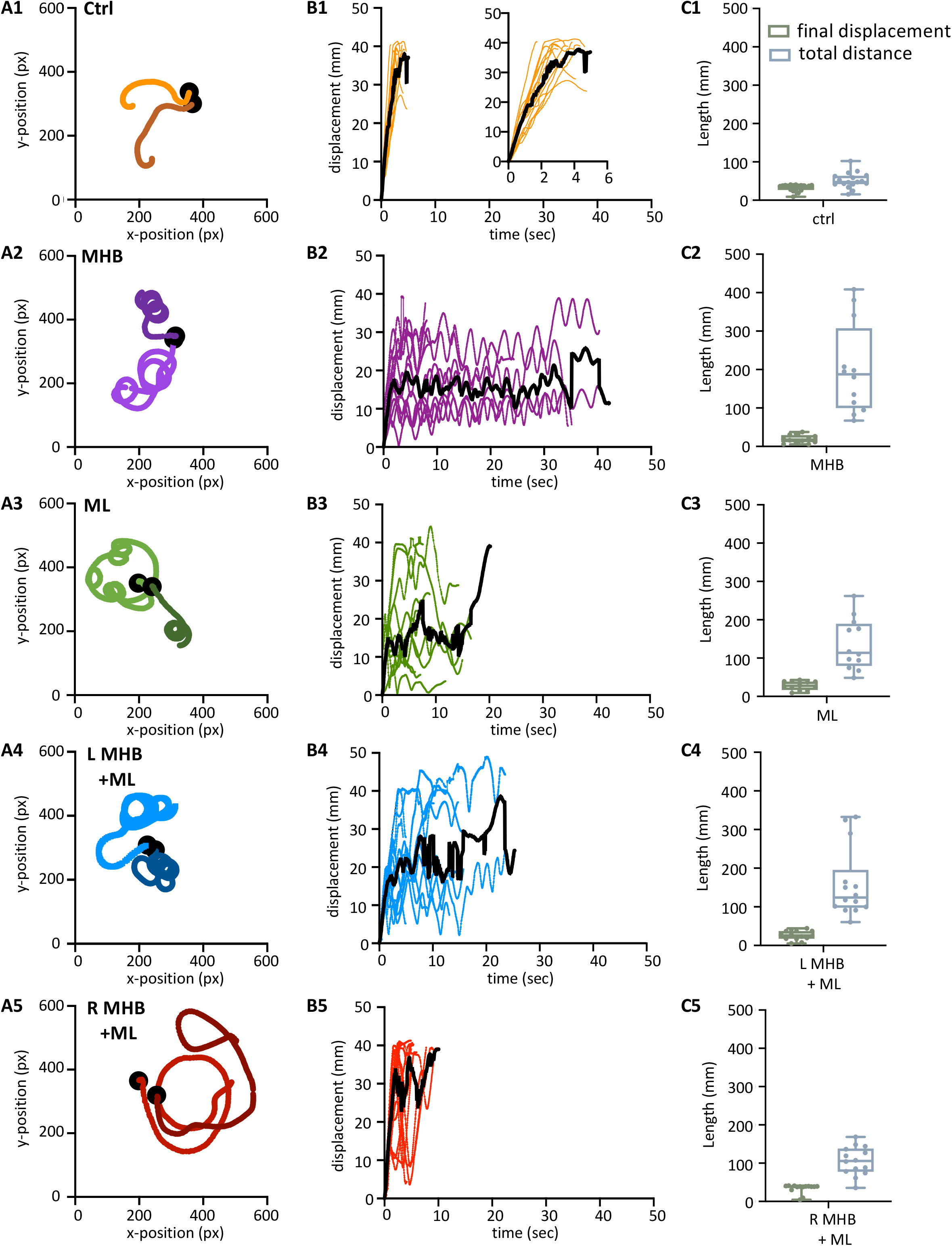
Midbrain lesions lead to stark changes in the trajectory of tadpole swimming. **(A)** Example swim trajectories (n=2) for each animal group extracted from the FastTrack Software. The starting position is symbolised by the black dot. For Panels A, B: Control: n=5, trials=12; MHB: n=6, trials=13; ML: n=4, trials=8; L MHB+ML: n=5, trials=10; R MHB+ML: n=4, trials=8. **(B)** Displacement time graphs of each animal trial (coloured lines) to show the tadpoles deviation from the starting position for each animal group (See Materials and Methods for calculation). The average is represented by the black line for each animals group. An increase in displacement refers to the tadpole swimming away from the starting position, while a decrease represents the tadpole swimming closer to the starting position. **(C)** Boxplot of the final displacement and total distance travelled for each animal trial within the respective animal group. For all panels, Control: n=10, trials=17; MHB: n=9, trials=12; Midline: n=8, trials=12; L MHB+ML: n=9, trials=14; R MHB+ML: n=10, trials=14. For boxplot, single data points are plotted, the middle horizontal line represents median latency, the box represents the interquartile range (IQR, 25-75 percentile), and the extended vertical bars represent the minimum and maximum values.

Swim trajectories were further analysed quantitatively by calculating the displacement for each tadpole per video frame (Figure 5B; see Materials and Methods for details). The displacement indicates the deviation of the tadpole’s swim position from the starting point (centre of the arena; 0 s) in each video frame up to the final frame when tadpoles ceased swimming (Figure 1J). In control animals (Figure 5B1; n=10 tadpoles, 17 trials), the steep increase in displacement over a relatively short period of time represents the forward direction of swimming as the tadpole swims further away from its starting position. Control animals showed an average peak of displacement at 37.9 mm (black line in Figure 5B1) due to the physical dimension of the arena used, which had a radius of 45 mm. Indeed, control tadpoles took mostly forward trajectories, quickly reached the wall of the Petri dish and stopped by contact and pressure applied to the head. On the other hand, the displacement of midbrain-lesioned animals, swimming predominantly in a circular pattern, was characterised by peaks and troughs indicative of repetitive circular movement of the animal throughout the entire swim cycle (Figure 5B2-5). MHB-(n=9 tadpoles, 12 trials) ML-(n=8 tadpoles, 12 trials) and L MHB + ML-lesioned animals (n=9 tadpoles, n=14 trials) adopted a consistent circular-shaped trajectory (Figure 5B2-4), while R MHB + ML-lesioned tadpoles (n=10 tadpoles, n=14 trials) had a more irregular trajectory, characterised by a brief forward start and wider circular movements (Figure 5B5).

Figure 5C summarizes these observations by presenting the median final displacement and total distance travelled for each experimental group (Figure 5C). The median values for displacement and total distance travelled by control animals are almost equal (Figure 5C1), as the control tadpole would constantly move away from the starting point, in an almost straight trajectory (Figure 5A1, B1). In contrast, midbrain-lesioned animals showed very different values for final displacement and total distance travelled (Figure 5C2-5). In fact, midbrain-lesion tadpoles covered longer distances, however due to the circular trajectories their final displacement values are smaller, because their swim episode ended nearer the starting position.

The above observations (Figure 5) also highlight that the midbrain lesions, despite altering the tadpole’s swim trajectory, still permit the animal to swim for a considerable amount of time and spontaneously stop. The limited size of the swim arena does not permit us to make direct comparisons of the duration of swim episodes between control and midbrain-lesioned animals, because in some trials tadpoles were forced to stop swimming through contact with the Petri dish walls. However, we further investigated the effect of midbrain lesions on the total duration of swim episodes, in all experimental groups, by taking into consideration the way each animal stopped swimming in each trial (spontaneous *vs* wall stops; Figure 6; number of tadpoles and trials per experimental group as mentioned above and in Figure 5). These data will also provide us with insights into possible changes to the tadpole’s stopping mechanisms due to midbrain lesions.

**FIGURE 6.**
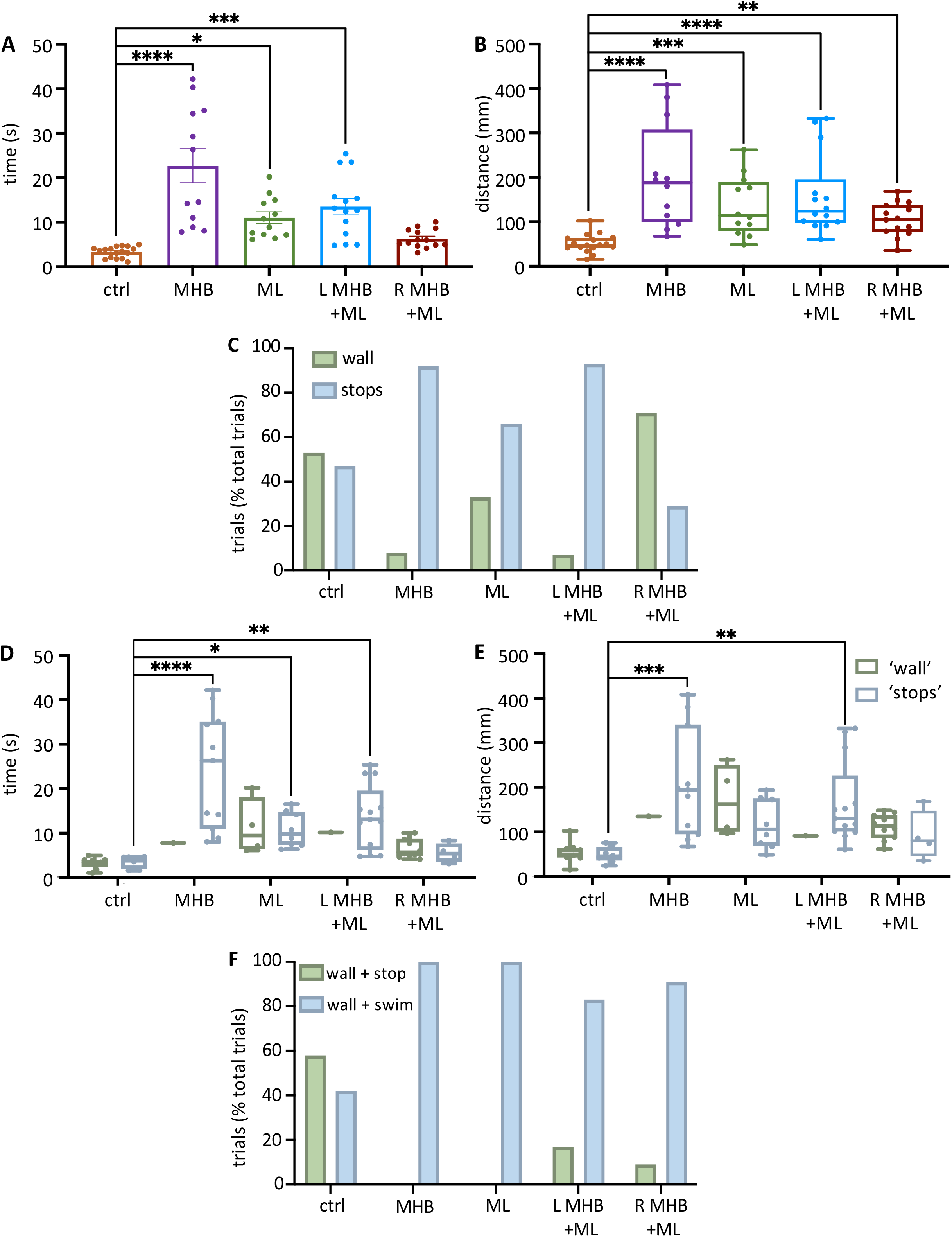
Midbrain lesions affect sustained swimming. **(A)** The average duration of swim episodes in control and midbrain lesioned animals. Controls swam for 3.3 ± 0.3 s after stimulation. MHB: 22.7 ± 3.8 s; ML: 11.0 ± 1.3 s; LMHB + ML: 13.5 ± 1.9 s; R MHB +ML: 6.3 ± 0.6 s. Reported as mean ± SEM. Single data points are plotted; the middle horizontal line represents the mean and the extended vertical bars represent the SEM. **(B)** Controls travelled 47.4, 41.9-63.7 mm (note the size of the arena had a radius of 45 mm). MHB: 187.5, 99.7-307.5 mm; ML: 113.7, 79.4-189.6 mm; L MHB +ML: 123.9, 97.7-195.7 mm; R MHB +ML: 105.8, 77.4-138.3 mm. **(C)** Tadpoles could stop either by hitting the edge of the arena (‘wall’) or stopping spontaneously (‘stops’). Control: 53% ‘wall’ vs 47% ‘stops’; MHB: 8% ‘wall’ vs 92% ‘stops’; ML: 33% ‘wall’ vs 66% ‘stops’; L MHB +ML 7% ‘wall’ vs 93% ‘stops’; R MHB +ML: 71% ‘wall’ vs 29% ‘stops’. **(D)** Boxplot for animals that stopped on the ‘wall’ or spontaneously (‘stops’). Control: 3.8, 1.9-4.6 s; MHB: 26.3, 11.0-35.1 s; ML: 9.8, 7.4-14.8 s; L MHB +ML: 13.1, 6.2-19.6 s; R MHB +ML: 5.4, 3.6-7.7 s **(E)** Distance swam for animals that stopped on the ‘wall’ or spontaneously (‘stops’). Control: 45.3, 35.8-66.6 mm; MHB: 194.6, 94.9-340.9 mm; ML: 105.7, 69.1-175.6 mm; L MHB +ML: 130.1, 100.7-227.1 mm; R MHB +ML: 79.9, 45.1-147.8 mm. **(F)** Percentage frequency for animals that stopped completely when obstructed by solid objects ‘wall + stop’ or carried on swimming ‘wall + swim’. For ‘wall + swim’; ctrl: 58% in 12 trials, MHB: 100% in 2 trials, ML: 100% in 6 trials, L MHB +ML: 83% in 6 trials, RMHB +ML: 91% in 11 trials. Sample size identical to those in fig. 6. For panels B, D and E, results are reported as median and IQR (25-75 percentile); for boxplot, single data points are plotted, the middle horizontal line represents median latency, the box represents the interquartile range (IQR, 25-75 percentile), and the extended vertical bars represent the minimum and maximum values. ****p<0.0001, ***p<0.001, **p<0.01 and *p<0.05 for Kruskal-Wallis/Dunn’s test and ANOVA/Dunnett’s

### The midbrain contributes to the tadpole’s stopping response

Figure 6A shows all data on duration of swim episodes irrespective of the way each animal’s swimming ceased. Control animals swam for 3.3±0.3 seconds (mean±SEM; s) after trunk skin stimulation (Figure 6A). The most significant increase in swim duration was seen in animals with complete disconnection of the midbrain from the rest of the brainstem (MHB; p<0.0001, ANOVA/Dunnett’s; MHB: 22.7±3.8 s). The ML and L MHB + ML lesions also led to overall significantly longer swim episodes, compared to control animals (ML: 10.99±1.32 s; p=0.0127, ANOVA/Dunnett’s and p=0.0003, ANOVA/Dunnett’s, L MHB + ML: 13.5±1.9 s). Consequently, the distance travelled will also increase in lesioned tadpoles (Figure 6B). Control tadpoles travelled 47.4, 41.9-63.7 mm (median, IQR), though the size of the arena (radius 45 mm) is a limiting factor here. MHB lesioned animals travelled the greatest distance of 187.5, 99.7-307.5 mm (median, IQR; p<0.0001, Kruskal-Wallis/Dunn’s test). Similarly, ML, L MHB + ML and R MHB + ML animals all travelled various distances than control (p=0.0005, Kruskal-Wallis/Dunn’s test; ML median: 113.7, IQR: 79.4-189.6 mm; p<0.0001, Kruskal-Wallis/Dunn’s test; L MHB +ML median: 123.9, IQR: 97.7-195.7 mm; p=0.0070, Kruskal-Wallis/Dunn’s test; R MHB +ML median: 105.8, IQR: 77.4-138.3 mm).

To address the limitations imposed by the arena size, data from each animal were categorised based on how the animal stopped swimming (Figure 6C). In this experimental setup, tadpoles can stop swimming either by hitting the edge of the arena (‘wall’) (Perrins et al., 2002;Li et al., 2003;Roberts et al., 2010) or spontaneously (‘stops’). Control animals stopped swimming at almost equal proportions by either bumping onto the walls of the arena or stopping spontaneously (53% ‘wall’ *vs* 47% ‘stops’ in 17 trials). In contrast the MHB, ML and L MHB + ML animals were less likely to encounter the walls of the Petri dish (Figure 6C; MHB: 8% ‘wall’ *vs* 92% ‘stops’ in 12 trials; ML: 33% ‘wall’ vs 66% ‘stops’ in 12 trials; L MHB + ML: 7% ‘wall’ vs 93% ‘stops’ in 14 trials), because their swimming had a circular trajectory. The clear majority of MHB tadpoles stopped spontaneously (8% ‘wall’ *vs* 92% ‘stops’ in 12 trials). The R MHB + ML tadpoles also swam in circular trajectories, however, those appeared to be wider when qualitatively compared to the circular trajectories of the other midbrain-lesioned tadpole groups (Figure 5A2-5). Thus, it is not surprising that the majority of the R MHB + ML tadpoles stopped swimming by bumping their heads onto the arena walls (71% ‘wall’ *vs* 29% ‘stops’ in 14 trials; Figure 6C).

Subsequent data analysis on the duration (time, s) and distance (mm) of swimming revealed that midbrain lesions significantly altered those swim parameters (Figure 6D, E). In trials where swimming stopped spontaneously (‘stops’), MHB (control median: 3.8, IQR: 1.9-4.7 s *vs* MHB median: 26.3, IQR: 11.0-35.1 s; p<0.0001, Kruskal-Wallis/Dunn’s test), ML (control median: 3.8, IQR: 1.9-4.7 s *vs* ML median: 9.8, IQR: 7.4-14.8 s; p=0.0154, Kruskal-Wallis/Dunn’s test) and L MHB + ML tadpoles (p=0.0011, Kruskal-Wallis/Dunn’s test; 13.1, 6.2-19.6 s) swam for significantly longer duration compared to control tadpoles (Figure 6D). This increase in duration of swimming was also followed by increase in the distance (mm) covered. More specifically, MHB (control median: 45.3, IQR: 35.8-66.6 mm *vs* MHB median: 194.6, IQR: 94.9-340.9 mm; p=0.0001, Kruskal-Wallis/Dunn’s test) and L MHB + ML animals (control median: 45.3, IQR: 35.8-66.6 mm *vs* L MHB + ML median: 130.1, IQR: 100.7-227.1 mm; p=0.0015, Kruskal-Wallis/Dunn’s test) swam the longest distance before stopping spontaneously (Figure 6E). No significant changes were detected in these aspects of swimming when tadpoles stopped due to contact with a physical object (Figure 6D, E). It is also worth noting that the limited size of the swim arena and the wider circular trajectories of the R MHB + ML tadpoles (Figure 5A5) enable these animals to reach the edge of the Petri dish, and this might partially contribute to the non-significant changes reported in Figure 6D, E (*vs* control tadpoles).

In our current experimental setup, animals did not always stop swimming when in contact with an object (wall of the Petri dish or Sylgard platform in the centre of the dish, Figure 1E), referred to as ‘wall + swim’ when tadpoles carried on swimming *vs* ‘wall + stop’ when tadpoles stopped after contact (Figure 6F). 58% of the trials performed on control tadpoles (n=10 tadpoles, 17 trials) led to the stopping of movement in response to head contact with a solid object. Midbrain lesions had stark effect on this behavior and ability of lesioned animals to stop movement in response to head contact with a solid object. In comparison to control animals, none of the MHB-(n=9 tadpoles, 12 trials) and ML-lesioned tadpoles (n=8 tadpoles, 12 trials) were able to stop swimming when their head encountered the object (Figure 6F; 100% of trials in MHB and ML tadpoles led to ‘wall + swim’). Similarly, the other two types of midbrain lesion also affected considerably the animals’ stopping mechanism (L MHB + ML: 83% wall + swim in n=9 tadpoles, 14 trials; R MHB + ML: 91% wall + swim in n=10 tadpoles, 14 trials; Figure 6F).

### The midbrain contributes to the postural orientation of the tail

It has been shown that in larva zebrafish excitability in the midbrain nMLF (nucleus of the medial longitudinal fasciculus) provides postural control for tail orientation, which in turn influences the direction (via steering) of swimming (Thiele et al., 2014). We investigated if the midbrain activity of the hatchling tadpole is essential in postural control of the tail, which in part might explain the changes seen in the swim trajectory of midbrain-lesioned animals. Tail deflection was measured as described in Methods and Figure 1F, while the tadpole was at rest, positioned into the Sylgard block groove with dorsal side up and the tail unrestrained, off the Sylgard block. Overall, all midbrain lesions resulted in increase in tail deviation in comparison to control tail position (Figure 7A, B1-5), however no statistical significance was recorded between the control mean tail angle and those of each lesioned group (p>0.05, ANOVA/Dunnett’s; Figure 7A). The unrestrained tail position of control animals (n=5, 12 trials), at rest, deviated by 0.9° ± 1.1° (mean ± SEM) with the greatest angle to the left at 8.2° and right at 5.1° (Figure 7A, B1). The R MHB + ML tadpoles showed the most bias in the position of the tail with a mean deviation angle of 7.8° ± 13.3° (mean ± SEM; n=4 tadpoles, 8 trials), and the largest angle to the left and right at 61.1° and 35.6°, respectively (Figure A. B5).

**FIGURE 7.**
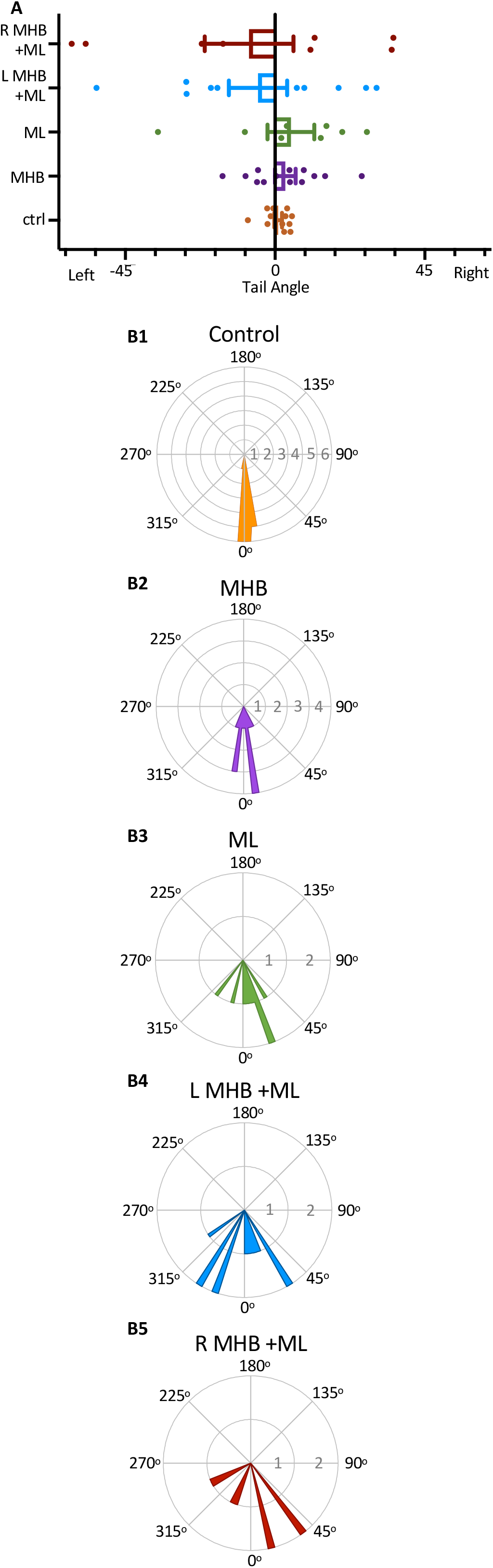
Midbrain lesions cause tail deflections. **(A)** The mean tail angle before at rest, prior to trunk skin stimulation. Ctrl: 0.9° ± 1.1°, n = 5 tadpoles, 12 trials; MHB: 3.1° ± 3.1°, n = 6, 13 trials; ML: 4.8° ± 7.0°, n = 4, 8 trials; L MHB+ML: -5.1° ± 8.7°, n = 5, 10 trials; R MHB+ML: -7.8° ± 13.3°, n = 4, 8 trials. Reported as mean ± SEM. Single data points are plotted, the middle horizontal line represents the mean, and the extended vertical bars represent the SEM. No significant difference between control and lesioned animals was identified. p>0.05 ANOVA/Dunnett’s. **(B)** Circular plot of the frequency distribution of the tail angle before swim starts. Each circular segment represents the number of values. Bin width is 5.8°.

## Discussion

The midbrain is an integral part of the supraspinal motor control network, shown to command locomotor activity from basal vertebrates to mammals (Shik et al., 1966;Shik et al., 1969;Sirota et al., 2000;Cabelguen et al., 2003;Uematsu et al., 2007;Dubuc et al., 2008;Jordan et al., 2008;Smetana et al., 2010;Ryczko and Dubuc, 2013;Daghfous et al., 2016). However, the midbrain neuronal circuitry and its connections with other supraspinal and spinal motor circuits are not fully understood. In this study we sought to investigate the role of midbrain in the control of coordinated locomotor behavior in the *Xenopus laevis* tadpole, a simple animal at a very early stage of development.

In the hatchling tadpole (developmental stage 37/38), the first evidence on the exact role of midbrain in the initiation of swimming (locomotion) came from studies on the pineal eye, situated above the midbrain-forebrain border (Foster and Roberts, 1982;Jamieson, 1997;Jamieson and Roberts, 1999). Light dimming excites pineal eye photoreceptors and axons of pineal ganglion cells project caudally into the ventral midbrain, without reaching the hindbrain and spinal cord. HRP (horseradish peroxidase) application in the hindbrain and spinal cord, discovered a cluster of midbrain neurons, so-called diencephalic/mesencephalic descending (D/MD) neurons (Jamieson and Roberts, 1999). The D/MD neurons were in a position where they could be contacted by the pineal ganglion cell axons and their activity was similar to that of the pineal. Furthermore, the axonal projections of the D/MD neurons to hindbrain and spinal cord (Jamieson and Roberts, 1999) indicate that these midbrain neurons could make direct ipsilateral connections and excite the central pattern generator (CPG) network. Midbrain neuron – CPG connections remain unidentified in the tadpole.

There is also evidence to support the involvement of the tadpole midbrain in the initiation and maintenance of swimming in response to trunk skin stimulation. Axons of the trunk skin sensory Rohon-Beard (RB) neurons excite dorsolateral ascending (dla) and dorsolateral commissural (dlc) sensory pathway neurons (Roberts et al., 2010;Roberts et al., 2019), whose axons have been shown to also reach the midbrain (Li et al., 2001). This suggests that the midbrain might play a significant role in the trunk skin sensorimotor pathway. However, connections from the sensory pathway to midbrain neurons remain unknown. The sensory pathway neurons (dla and dlc) initiate or accelerate swimming by amplifying excitation (Li et al., 2001;Li et al., 2004b), however, their contribution to the rostral CNS does not appear to influence the generation of tadpole swimming. The hatchling tadpole can still generate episodes of sustained swimming after removal of the rostral CNS (Li et al., 2006), as well as, the disconnection of the midbrain from the rest of the brainstem and spinal cord. We also confirm this here, in both behavioral and electrophysiological experiments transection along the midbrain-hindbrain (MHB) border does not prevent the generation of swimming *per se*. So, what might be the role of the midbrain within the trunk skin sensorimotor pathway?

To answer this question, we used a combination of midbrain lesions and trunk skin stimulation, in behavioral and electrophysiological settings, to identify the specific consequences of each midbrain lesion on kinematic parameters associated with the hatchling tadpole swimming. We reveal that the tadpole relies heavily on the midbrain for the timely and effective initiation, as well as arrest of swimming.

### Timely and coordinated initiation of swimming is dependent on the midbrain

Our behavioral experiments show that following trunk skin stimulation control tadpoles initiate swimming at variable latencies and with preference for a contralateral start, which agrees with previous studies (Buhl et al., 2015;Koutsikou et al., 2018;Ferrario et al., 2021;Messa, 2022). Midbrain lesions affected both of those parameters to varying degrees, without preventing the generation and maintenance of locomotion. This agrees with studies on salamander and lamprey, at later stages of development, where just unilateral excitation of the MLR could lead to bilateral activation of reticulospinal neurons (Ryczko et al., 2016). The MHB (Midbrain/Hindbrain Border) and the R MHB + ML (Right side of the MHB + Midline) lesions led to significant increases in the swim latency when the left side of the trunk skin was stimulated. First, this is in accordance with tadpole studies that identified a population of caudal hindbrain descending interneurons (hdINs) being sufficient for driving swimming in this animal even in the absence of the rostral brainstem (Li et al., 2006;Soffe et al., 2009). Second, studies have demonstrated that the contralateral to the stimulus side, which in our experiments is lesioned in MHB and R MHB + ML tadpoles, is the fastest and strongest (Jamieson and Roberts, 1999;Buhl et al., 2015). In support, the ML and L MHB + ML (ipsilateral to the stimulus) lesions, did not lead to significant increases in latency, most likely because in these lesions and unlike in the ones of MHB and R MHB + ML, sensory information, especially via the dlc pathway, could still reach the midbrain on the side contralateral to the stimulus and permit top-down descending control to be exerted by the fastest, contralateral pathway.

The behavioral results also indicate that disruption of input and output information from the contralateral midbrain, here in ML and R MHB + ML lesions, had stark effects on the side of swim initiation. Tadpoles with these lesions mostly responded to trunk skin stimulation with ipsilateral initiations (here the intact side of the CNS). This suggests that the blockade of the strongest, dlc pathway carrying sensory information to the contralateral midbrain together with the lack of *(i)* communication between the two sides of the midbrain and *(ii)* descending control from the contralateral midbrain force the animal to ipsilateral motor bias. Such functional lateralization and asymmetric motor behaviors have been seen in many species (Rogers, 2009), however, the underlying neural substrates and connecting pathways remain unknown. Recently, identification of an intrinsic lateralized behavior in zebrafish larvae showed that the left/right motor bias is dependent on neurons in the diencephalon that project to the habenula (Horstick et al., 2020). The rest of the lesions utilized here caused minor changes to the ratio of contralateral *vs* ipsilateral starts, thus sustaining the contralateral motor bias. Overall, these data highlight that even at this early stage of development, the midbrain can modulate the side and latency of swim initiation in response to trunk skin stimulation, which makes it essential to the timely and efficient initiation of this behavioral motor response.

The significant changes in latency to swim initiation following MHB lesions were further investigated in experiments of fictive swimming in hatchling tadpoles. As soon as swimming is initiated, the neuronal activity of hindbrain and spinal cord CPG alternates rhythmically between the two side via reciprocal inhibition, which will be maintained duing ongoing swimming (Roberts et al., 2010;Moult et al., 2013). Here we show that MHB-lesioned tadpoles exhibit synchronous oscillations at initiation, referred to as ‘synchrony’, a phenomenon that has been previously observed in the hatchling tadpole (Kahn and Roberts, 1982;Li et al., 2014). Simultaneous ventral root (VR) activity on opposite sides of the trunk is indicative of immobility until the animal can engage in an antiphase oscillatory muscle activity, which in a behavioral setting can lead to full body propulsion. Our data suggest that in the hatchling tadpole midbrain descending control is necessary for the generation of antiphasic activity patterns by contributing to the avoidance of synchrony during swim initiation. However, the synchronous VR events recorded from MHB animals do not fully explain the increase in latency to swim initiation observed in behavioral experiments. This is due to threshold stimulation of MHB tadpoles leading to shorter latencies despite the increase in synchronous events. It is unknown how the tadpole midbrain interacts with the brainstem and spinal cord to avoid synchronous initiations. We suggest that the midbrain might directly or indirectly promote downstream unilateral inhibition, which limits the firing of motoneurons and other CPG interneurons (Li et al., 2004a;Koyama et al., 2016;Liu and Hale, 2017;Koyama and Pujala, 2018). This explanation is also in line with studies showing that blockade of glycinergic inhibition leads to synchrony in neonatal rats (Cowley and Schmidt, 1995) and lamprey (Cohen and Harris-Warrick, 1984).

### The midbrain is essential for the discrimination of stimulus strength

Shik and colleagues (1966, 1969) demonstrated, on the decerebrate cat, that MLR stimulation at different strengths produces different motor behaviors. Low-strength stimulation produced walking, while an increase in the stimulus strength led to trot and gallop type movements. The increase in stimulus strength leads to increase in recruitment of active MLR neurons. Similarly, we show in the young *Xenopus* that different stimulus strengths lead to distinct motor outcomes with the midbrain being essential for this sensory discrimination. The hatchling tadpole can discriminate stimulus saliency, because electrical stimulation at different intensities (threshold *vs* suprathreshold) led to distinct patterns of swim initiation in control tadpoles. Suprathreshold stimulation shortened the median swim initiation latency of control animals to more than half, in comparison to threshold stimulation. In addition, the stronger stimulus led to a contralateral motor bias with a swim latency significantly shorter in comparison to the ipsilateral motor response latency (in the fewer occasions when the ipsilateral side started first). This difference between ipsilateral and contralateral latencies is not present in control tadpoles stimulated at threshold intensities. These data also agree with previously published results, that show suprathreshold stimulation leads to contralateral motor initiation bias, with no overlapping latencies between ipsilateral and contralateral motor responses (Zhao et al., 1998). The authors showed very short delay glycinergic IPSPs in motoneurons ipsilaterally to stimulation, whilst no inhibition was found in motoneurons on the contralateral side (Zhao et al., 1998). Interestingly, this glycinergic inhibition was reported only at higher strength stimuli and no IPSPs were recorded following threshold stimulation (Zhao et al., 1998). Based on the short latency of these IPSPs the authors proposed that ascending interneurons (aINs) might be responsible for the ipsilateral inhibition prior to the initiation of swimming (Zhao et al., 1998), which implies that dlas (the ipsilateral sensory pathway neurons) form synaptic contacts with aINs. This connect could explain how aINs would be responsible for the short-delayed inhibition recorded in ipsilateral motoneurons (Zhao et al., 1998). Our data here do not provide the underlying mechanistic differences, however from an ethological point of view it is advantageous for the animal to move away from the source of a stimulus that can be potentially threatening to its survival.

The MHB lesion prevented all animals from maintaining the discrimination between different strengths of stimulation. The lesioned tadpoles responded to either stimulus (threshold vs suprathreshold) with similar latency, which differs from data from control animals responding with a faster swim initiation to suprathreshold stimulus.

### Midbrain control of posture and swim trajectory

The variable change or loss of behavioral motor functions of the hatchling tadpole, due to distinct midbrain lesions, also include the swim trajectory, displacement, duration, and distance, as well as arrest of swimming. Our data show that any type of ablation in the midbrain results in a deflection of the tail. This suggests that the midbrain is a source of activity before any external stimulus is applied and locomotion is initiated in response to that. However, the midbrain neurons responsible for this activity have not been identified. Our findings are consistent with previous literature in zebrafish, demonstrating that ablation of midbrain nMLF neurons also causes pronounced deflections of the tail (Gahtan et al., 2005;Thiele et al., 2014). We propose that the changes observed in the swim kinematics mentioned above are due to tail deflection, which changes the yaw angle between the head and the tail resulting in loss of postural control that cannot be sustained even after partial loss of midbrain descending control.

We also provide strong evidence that tadpole midbrain activity is implicated in the stopping of swimming following contact of the head with an object, in this case the wall of the Petri dish. Previous studies have shown that when the tadpole head skin is pressed, trigeminal ganglion sensory neurons are activated, which in turn project to the hindbrain (Roberts and Blight, 1975;Roberts, 1980). In the hindbrain the trigeminal sensory neurons target the so-called midhindbrain reticulospinal neurons (mhrs), which produce GABA-mediated inhibition of neurons active in swimming, while mhrs are inhibited during sustained swimming (Roberts et al., 1987;Perrins et al., 2002;Li et al., 2003). Recent work investigating the swim stopping mechanism in the hatchling tadpole revealed the involvement of a cholinergic pathway which via muscarinic M2 receptors leads to opening of G protein-coupled inward-rectifying potassium channels (GIRKs), which in turn inhibit CPG neurons essential to swim initiation and maintenance (Li et al., 2017). The exact interactions of the tadpole midbrain with the rest of the brainstem stopping mechanism remain unknown. However, our results are in line with previous evidence from experimental work on higher vertebrates, showing that the command for locomotion arrest is integrated in supraspinal centres (Takakusaki et al., 2003;Bouvier et al., 2015).

### Identity of midbrain neurons involved in the trunk skin sensorimotor pathway

The light dimming sensorimotor pathway, from the pineal eye to D/MD neurons to the brainstem and spinal CPG and motoneurons exhibits latencies from light dim to fictive swimming of 70 – 110 ms (Roberts, 1990). This range of onset latencies is very similar to the fictive swim latencies following trunk skin stimulation (Koutsikou et al., 2018). So the similarity in onset latencies between the two sensorimotor pathways indicates that the D/MD neurons might also be involved in the trunk skin sensorimotor pathway as third order neurons (Jamieson and Roberts, 1999), following synaptic connections from dla and dlc sensory pathway neurons which are activated by RB cells (Roberts et al., 2010) and whose axons can reach the midbrain (Li et al., 2001).

Although we cannot exclude the possibility that D/MD neuronal excitability contributes to trunk skin-evoked swimming, their contribution alone does not fully explain our findings. The ML (midbrain midline) lesion used in this study severs commissural connections between the two sides of the midbrain and leads to significant changes in kinematics of swim behavior in response to trunk skin stimulation. Previous data have shown that the identified D/MD neurons involved in the light dim pathway, do not have commissural axons (Jamieson and Roberts, 1999) and the dlc neurons that carry sensory information from the trunk skin to the brainstem do not cross the midline at the level of the midbrain (Roberts et al., 2010). Thus, we propose that the effects of ML lesion seen here are only possible if there is another group(s) of midbrain neurons that *(i)* respond to trunk skin excitation, via synaptic connections with dla and dlc neurons and *(ii)* can transfer this excitation to the opposite side at the level of the midbrain. Interestingly, HRP backfill experiments by Jamieson and Roberts (1999) identified the D/MD group in a cluster close to the midbrain-forebrain border, where these neurons could be contacted by the axons of the pineal ganglion cells (Foster and Roberts, 1982;Jamieson and Roberts, 1999). In the Jamieson and Roberts (1999) study the location of the HRP retrograde application in the hindbrain is defined as rostral to the 5^th^ postotic myotome. It is possible that the authors did not fully identify the population(s) of midbrain neurons that could be activated by trunk skin stimulation and have commissural axons that also descend to the hindbrain because they restricted the area of HRP application. Based on the similarities of onset latencies described above, it is possible that the trunk skin-activated midbrain neurons might synapse onto rostral (pre-otic) hindbrain extension neurons (exNs with command properties; (Messa and Koutsikou, 2021)), whose recurrent excitatory population produces the relatively slow and variable onset to swimming in the tadpole (Koutsikou et al., 2018;Roberts et al., 2019;Ferrario et al., 2021). Further work is required to both identify the connections between midbrain neurons activated by trunk skin stimulation and their synaptic connections to exNs and supraspinal CPG.

### Concluding comments

Like all animals, the hatchling *Xenopus* tadpole needs to be able to navigate its environment and avoid predators. We identified that the midbrain of this animal contributes significantly to its survival by contributing to postural control, as well as the timely and efficient swim initiation, maintenance, and arrest in response to external stimuli. Thus, this animal’s primitive midbrain neurons share functions of sensorimotor descending control with midbrain and brainstem neurons in older and higher vertebrates. This makes the tadpole, with its very simple CNS, an ideal animal model to investigate the underlying circuitry of supraspinal sensorimotor control at the single cell level.

## Acknowledgments

The authors would like to acknowledge the financial support of The Physiological Society UK by awarding a research grant to SK.

## Notes

### Competing Interest Statement

The authors have declared no competing interest.

